# Number and proportion of P. *falciparum* gametocytes vary from acute infection to chronic parasite carriage despite unaltered sexual commitment rate

**DOI:** 10.1101/2021.11.05.467456

**Authors:** Hannah van Dijk, Martin Kampmann, Nathalia F Lima, Michael Gabel, Usama Dabbas, Safiatou Doumbo, Hamidou Cisse, Shanping Li, Myriam Jeninga, Richard Thomson-Luque, Didier Doumtabe, Michaela Petter, Kassoum Kayentao, Aissata Ongoiba, Teun Bousema, Peter D Crompton, Boubacar Traore, Frederik Graw, Silvia Portugal

## Abstract

*Plasmodium falciparum* infections persist through long dry seasons at low parasitaemia without causing malaria symptoms and thus remain untreated. In asymptomatic children, increased circulation of infected erythrocytes without adhering to the vascular endothelium is observed during the dry months, compared to febrile malaria in the wet season. However, alterations of parasite sexual commitment and gametocytogenesis have not been investigated. Here, we compared the expression of genes related to sexual commitment and gametocytogenesis, the proportion and density of *P. falciparum* gametocytes, and the blood concentration of phospholipids in dry season asymptomatic individuals versus symptomatic subjects in the wet season. Additionally, we adapted a within-host mathematical model considering asexual and sexually-committed parasites and gametocytes to understand the dynamics of gametocyte number and proportion as infections progress. Compared to clinical malaria cases, transcripts of late-stage gametocytes were predominantly upregulated in the dry season, associating with increased proportions of mature gametocytes; while transcription of genes related to parasite sexual commitment was unaltered throughout the year. Our data suggest that gametocyte density and proportion diverge as infections progress from recent transmission to chronic carriage, without alterations in the sexual commitment rate over time.

## Introduction

The malaria-causing parasite *Plasmodium falciparum* requires a mosquito to complete its lifecycle and be transmitted between humans^1^. During blood-feeding, infected mosquitoes deposit sporozoites in the human dermis that can reach the circulation, arrive at the liver and infect hepatocytes, where thousands of merozoites are formed and later released into the blood stream to start continuous asexual intra-erythrocytic replication cycles of ∼48h. A small proportion of asexual parasites commit to sexual development^2^. Instead of another ∼48h asexual replication leading to 8-30 new merozoites ready to continue asexual cycles, the parasite develops into a sexually-committed schizont producing the same number of sexually-committed merozoites. Upon infection of new red blood cells (RBCs) each sexually-committed merozoite produces one stage I gametocyte^3,4^ that in 8-12 days goes through five developmental stages^5^. Stages II-IV of gametocyte development occur in the bone marrow or spleen^6,7^, and therefore do not circulate; while stage V gametocytes circulate up to ∼20 days^8,9^, waiting for another mosquito-bite. In the mosquito midgut, gametocytes transform into female and male gametes that fuse into a zygote, which develops into an ookinete. In the midgut wall, the ookinete generates sporozoites that reach the mosquito salivary glands and can restart the lifecycle. Continuous presence of mosquitoes and humans allows perennial malaria transmission. However, in areas with months-long dry season, the mosquito population plummets to zero^10^, transmission is interrupted and *P. falciparum* survives in humans until mosquitos return in the ensuing wet season^11-13^. Long mosquito-free periods make gametocyte production a dead-end since these cannot survive to bridge two wet seasons^8,9^. Dynamically adapting sexual commitment rates depending on transmission environment, e.g. through sensing of dry-season associated cues, would represent a benefit for the parasite, as waste of resources could be avoided. Whether the parasite dynamically alters resources allocation into transmission to the mosquito to adapt to environmental changes remains unclear. Various factors such as host age^14^, haematocrit^15^, haemoglobin type^16^ and duration of symptoms^17^ have been linked to gametocytaemia in individuals with clinical malaria. Higher frequencies of gametocyte-positive individuals were observed in chronic asymptomatic infections compared to acute infections^18-22^. In vitro conditions have been shown to directly influence sexual commitment and gametocytogenesis^3,23-25^. Lysophosphatidylcholine (Lyso-PC) has been identified as repressor of the pro-sexual commitment transcription factor AP2G, and low levels of Lyso-PC have been found to increase the proportion of sexual forms in vitro^25^. Gametocyte development protein 1 (GDV1) was described as the de-repressing effector, which promotes *ap2g* expression and induces gametocytogenesis^26^, and homeodomain protein 1 (HDP1) acts as a positive regulator of gametocyte development^27^. A shift in intracellular S-adenosylmethionine/S-adenosyl homocysteine has been shown to affect histone methylation and increase parasites commitment to sexual differentiation^28^. Still, it remains unclear how sexual commitment rates are regulated in vivo and whether the parasite can optimize its transmission-investment. A study on controlled human malaria infections investigated whether mosquito bites, a plausible seasonal cue, could induce gametocytogenesis found no evidence of such an effect ^29^.

Dry season *P. falciparum* parasites exhibit a transcription pattern distinct of that of parasites causing febrile malaria during the wet season^11,30^. These seasonal transcriptional differences reflect longer parasite circulation within each ∼48h intra-erythrocytic cycle in the dry season, leading to increased splenic clearance of iRBCs and contributing to the low parasitaemias observed during the dry season^11^, but alterations of parasite sexual commitment and gametocytogenesis remain unexplored.

Here, we compared sexual commitment and gametocytogenesis of parasites persisting through the dry season with those of parasites in clinical malaria cases. We found that clinical malaria cases consistently exhibit higher parasite densities compared to asymptomatic infections, and that expression of marker genes of mature gametocytes and the proportion of gametocytes among all parasites were higher in persisting infections; while sexual commitment did not substantially vary longitudinally within each individual. Additionally, we determined that serological differences observed throughout the year were unable to differently induce *P. falciparum* gametocytogenesis in vitro. Finally, adapting a previously described within-host mathematical model^31^, we inferred the dynamics of gametocyte number and proportion as infections progress from recently transmitted through chronic asymptomatic dry season carriage. We conclude that a constant sexual commitment rate allows for dynamic alterations of sexual stages of *P. falciparum* consistent with empirical observations.

## RESULTS

### Seasonal transcriptional patterns of *P. falciparum* sexual development, and lipid concentration in plasma

Previously, we described transcriptional differences between *P. falciparum* from asymptomatic children at the end of the dry season and children with clinical malaria in the wet season^11^. Now, we extend our analysis to focus on sexual development across the two conditions and started by compiling a list of genes related to gametocytogenesis from four RNAseq published studies to query potential adaptations of *P. falciparum* sexual stages to asymptomatic persistence (Supplementary Table 1). Painter et al. reported 786 gametocyte-related transcripts in vitro^32^, of which we detected 333 in the dry vs wet season dataset^11^. Poran et al. described early and late gametocyte transcripts comparing day 2 vs 9 of gametocytogenesis^33^, we found that 2 out of 3 early gametocyte transcripts and 15 out of 17 late-gametocyte transcripts were detected in the dry vs wet season dataset. Lopez-Barragan et al. compared days 4 vs 15 of gametocytogenesis and identified 41 early- and 58 late-gametocyte genes^34^. Of those, 32 and 44 were detected in the dry vs wet season dataset^11^. Finally, Jeninga et al.^35^ compared days 4 vs 10 of gametocytogenesis and identified 237 transcripts assigned to early-gametocytes independent of sex, as well as 314 late gametocyte genes. Out of these, 80 early- and 133 late-gametocyte genes were detected at the end of dry vs wet season dataset^11^. The full list of 1258 sexual commitment- and gametocytogenesis-related genes showed limited overlap between reports (Supplementary Table 1), potentially due to the different timepoints and comparison conditions defined in the different studies. In summary, in the end of dry season asymptomatic vs wet season clinical cases RNAseq dataset^11^ we detected 551 of the 1258 genes, which could be expressed in circulating sexually-committed asexual stages, stage I or stage V gametocytes. Principal component analysis of expression values of these 551 genes expressed in *P. falciparum* of asymptomatic children at the end of the dry season and children with clinical malaria in the wet season^11^ revealed segregation of transcription profiles based on seasonality and clinical presentation (Fig. 1A), and clustering by similarity also produced visible seasonal patterns (Supplementary Fig. 1). We performed gene set enrichment analysis (GSEA) to assess whether gametocyte-associated genes (described by López-Barragán et al., Jeninga et al., Poran et al., and Painter et al.), were modulated in dry season infections. Genes from each study were analyzed separately, and López-Barragán et al. and Jeninga et al. also allowed distinguishing between early- and late-stage gametocytes.

**Figure 1.**
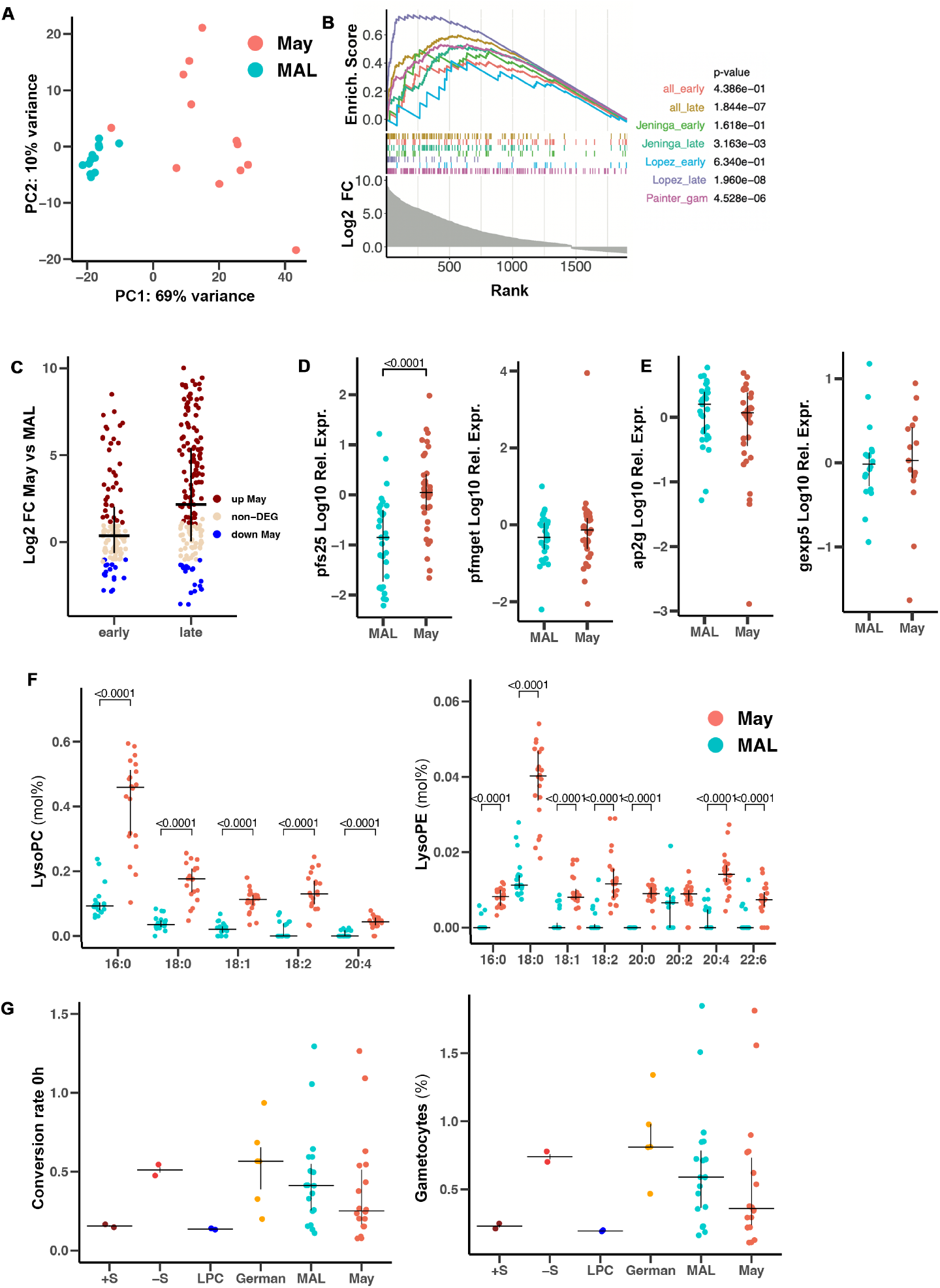
*P. falciparum* transcription and gametocytogenesis in clinical malaria and at the end of the dry season. **A** Principal components analysis of RNA-Seq data of 551 transcripts gametocyte specific *P. falciparum* genes from parasites collected at the end of the dry season and from clinical malaria cases (n= 12 May, 12 MAL). **B** Gene set enrichment analysis of gametocyte associated gene published by López-Barragán et al.; Jeninga et al., Painter et al. among DEGs between parasites collected at the end of the dry season (n=12) and from clinical malaria cases (n=12). All_early and All_late sets are collated from Lopez-Barragan et al., and Jeninga et al. and gametocyte associated genes published in Poran et al.. DEGs identified between clinical malaria cases and dry season asymptomatic infections were ranked by log2fold-change and GSEA-analysis performed with the fgsea algorithm. **C** Fold change of early and late gametocyte transcripts between parasites collected at the end of the dry season and from clinical malaria cases (n= 12 May, 12 MAL) according to López-Barragán et al.; Poran et al., and Jeninga et al.. Each dot represents a gene and color identifies fold change direction. **D** Relative expression of markers of female (*pfs25*, left) and male gametocytes (*pfmget*, right) to a housekeeping gene (*glycine-tRNA-ligase*) detected by qRT-PCR in clinical malaria cases (MAL, n=31) and in RDT^+^ subclinical children at the end of the dry season (May, n= 37). **E** Relative expression of *ap2g* and *gexp5* detected by qRT-PCR in clinical malaria cases and in RDT^+^ subclinical children at the end of the dry season. (MAL n=31, and May n=37 for *ap2g* and MAL n=18, and May n=15 for *gexp5*). **F** Plasma concentration of Lyso-PC (left) and Lyso-PE (right) species in RDT+ subclinical children at the end of the dry season (May, n=20) and in children with their first clinical malaria episode (MAL, n=20) in the wet season. **G** Pf2004 conversion rate (gametocytes d5/parasites d2) (left) and gametocyte density (right) detected by microscopy, after 24h culture with 25% plasma collected during clinical malaria episodes in the wet season (MAL n=19) or at the end of the dry season (May n=18), or German pooled plasma (German) alongside without serum (-S), serum supplemented (+S) and Lyso-PC supplemented (LPC) conditions. Lipid data and expression values represented as median ± IQR; Wilcoxon test between MAL and May with Bonferroni multiple comparison correction. Conversion rates represented as median ± IQR; Wilcoxon test.

Furthermore, we combined all early- and all late-stage defining genes into early and late gametocyte supersets. Due to the small number of genes reported by Poran et al., these data were not individually analyzed but incorporated into the all early and all late supersets. We identified statistically significant enrichment of the Jeninga et al. late-gametocyte gene set (normalized enrichment score, NES=1.29, p-value=4.053*10^-3), the López-Barragan et al. late-gametocyte gene set, (NES=1.76, p-value=9.235*10^-9), as well as the late gametocyte superset (NES=1.49, p-value=3.11*10^-7), and the Painter et al. gene set of gametocyte-associated genes (NES=1.42, p-value=5.01*10^-6) among genes up-regulated at the end of the dry season vs wet season clinical malaria cases, while none of the early-gametocyte gene sets afforded statistically significant enrichment (Fig. 1B). Accordingly, we observed a higher proportion of the late-gametocyte specific transcripts upregulated in the end of dry season asymptomatic vs wet season clinical cases RNAseq dataset^11^ (Supplementary Table 1) compared to the proportion of early-gametocyte specific transcripts. Close to two thirds (65%) of late-gametocyte transcripts were upregulated in the dry season (112/172 genes, Chi^2^ *p*-value < 0.00001), while only 39% (43/109 genes, Chi^2^ *p*-value=0.36784) of early-gametocyte transcripts, and 33% of total *P. falciparum* transcripts (1131/3381 genes) were upregulated at the end of the dry season in May (Fig. 1C). Markers of female (*pfs25*, PF3D7_1031000) and male (*pfmget*, Pf3D7_1469900) late-stage gametocytes were among the upregulated transcripts at the end of the dry season, in May, in the same RNAseq dataset^11^ (Supplementary Table 1).

These transcriptional differences prompted us to quantify the relative expression of female and male gametocyte-coding genes in 68 samples (MAL n=31, May n= 37) collected from the same cohort study as the reported RNAseq data^11^, from asymptomatically infected children at the end of the dry season (May) and age and sex-matched children presenting with clinical malaria symptoms in the wet season (MAL). qRT-PCR of marker genes of female (*pfs25*, PF3D7_1031000) and male (*pfmget*, Pf3D7_1469900) gametocytes relative to a housekeeping gene (*glycine-tRNA ligase*, PF3D7_1420400) confirmed the increased expression of the female gametocyte associated gene in May at the ebd of the dry season, compared to malaria cases (Fig. 1D). Conversely, transcripts known to regulate sexual commitment (*ap2g*, PF3D7_1222600; and *gdv1*, PF3D7_0935400), and transcripts expressed by sexually-committed schizonts (*msrp1*, PF3D7_1335000), or by early-gametocytes (*gexp5*, PF3D7_0936600) did not appear differentially expressed in the end of dry season asymptomatic vs wet season malaria cases RNAseq dataset^11^ (Supplementary Table 1). qRT-PCR analysis of the regulator of sexual commitment, *ap2g*, and the early gametocyte marker *gexp5* confirmed the absence of significant differences between the conditions (Fig. 1E). We then performed shotgun lipidomics with 40 samples from the same study site to examine potential associations between gametocyte proportion and plasma Lyso-PC and lysophosphatidylethanolamine (Lyso-PE) composition. The lower expression of late-gametocyte genes in the 20 malaria cases (MAL) coincided with lower levels of several species of the Lyso-PC and Lyso-PE families compared to the 20 asymptomatic carriers at end of the dry season (May) (Fig. 1F), suggesting that LysoPC levels might not be responsible for the observed gene expression changes, but rather a consequence of high parasitaemias and inflammation during acute infection.

Additionally, we interrogated whether the low phospholipids levels in plasma of malaria cases in the wet season compared to dry season persistent dry season infections in May, could sexual conversion and result in different gametocytaemia in vitro. We exposed *P. falciparum* of the Pf2004 strain to plasma collected from 19 clinical malaria patients in the wet season (MAL) or from 18 asymptomatically infected children at the end of the dry season (May), alongside the previously described conditions^25^ to induce or inhibit sexual conversion and gametocyteamia. As reported^25^, we observed that 24h after culture without serum (-S), and thus without Lyso-PC, sexual conversion rate increased compared to parasites cultured in the presence of 10% serum (+S) or without serum but supplemented with 20 µM Lyso-PC (LPC) (Fig. 1G). However, conversion rates into gametocytes five days after treatment did not show a statistically significant difference between supplementation with 25% of plasma from children with clinical malaria (MAL), plasma from dry season asymptomatic carriers (May), or German pooled plasma (German) (Fig. 1G). The induced sexual commitment rates were variable and did not associate to disease status or season of the tested plasma, indicating that the differences in Lyso-PC between asymptomatic children and malaria cases (Fig. 1F) were insufficient to impose alterations in sexual commitment or gametocytaemia.

### Sexual commitment and gametocytogenesis dynamic in persisting infections

Next, we obtained plasma and parasite samples collected longitudinally throughout the dry and wet seasons in the same cohort study, to investigated associations between expression of gametocyte-related genes and plasma Lyso-PC in children continuously infected from the wet until the end of the dry season, and in children with clinical malaria cases in the wet season. Within previously reported metabolomic data of this cohort ^36^, we found that Lyso-PC 18:0 was lower in clinical malaria cases (MAL n=15) compared to asymptomatic infections from different timepoints (Oct n=17, Mar n=19, and May n=19); and it also decreased from the wet (Oct) to the end of the dry season (May) (Fig. 2A). Lyso-PC 18:0 levels in persistent infections (Oct, Mar and May) were comparable to those in non-infected individuals at the end of the dry season (May^−^), indicating little or no impact of asymptomatic infections on Lyso-PC 18:0 (Fig. 2A).

**Figure 2.**
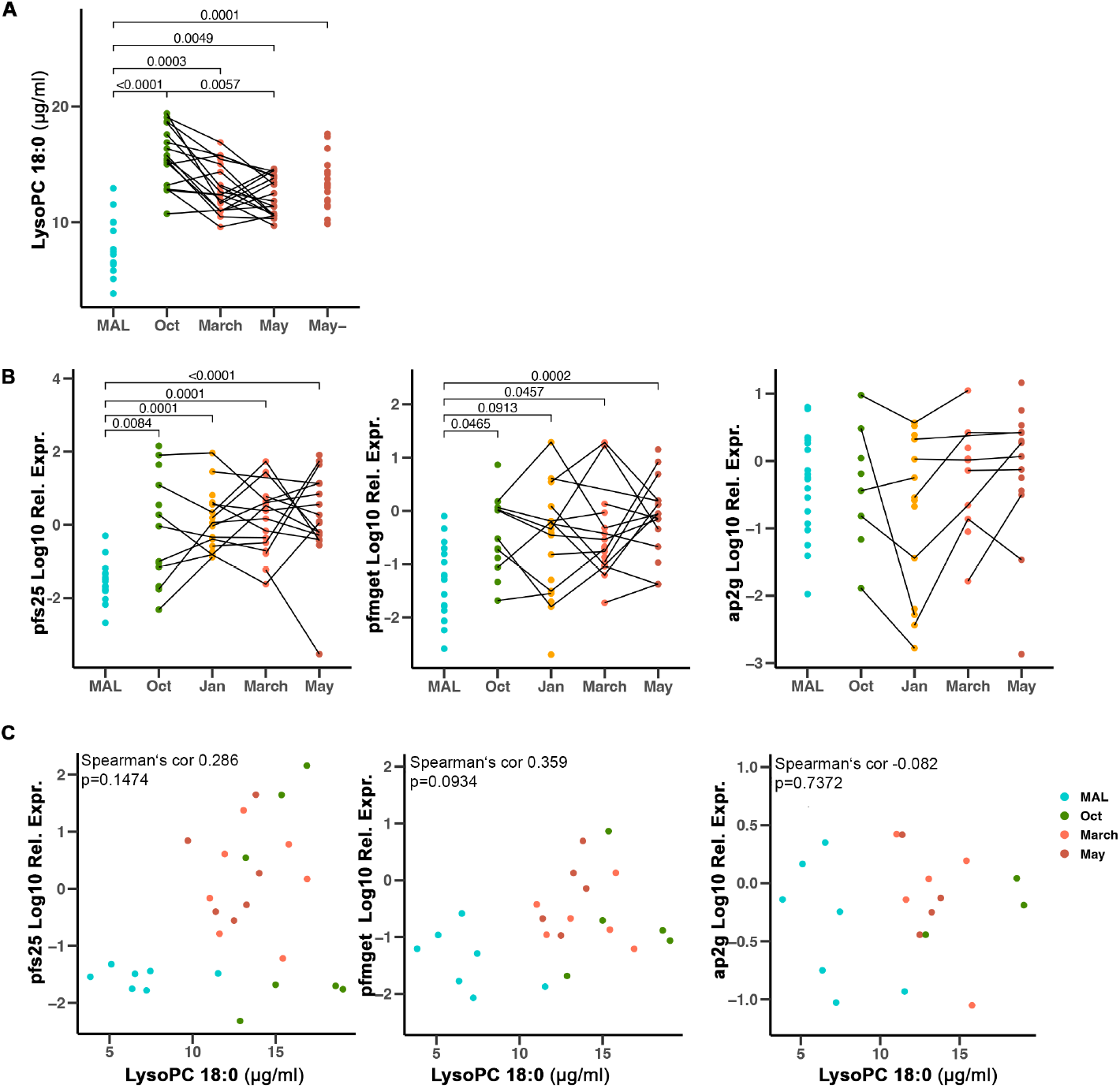
Plasma concentration of LysoPC and its effect on sexual commitment in persistent and clinical infections. **A** Plasma concentration of Lyso-PC 18:0 during clinical malaria episodes (MAL n=15), RDT^+^ subclinical infection times during the wet (Oct n=17) and the dry season (Mar and May n=19), and of uninfected children at the end of the dry season (May-n=20). **B** Relative expression of markers of female gametocytes (*pfs25*, left), male gametocytes (*pfmget*, centre), and sexual commitment (*ap2g*, right) relative to a housekeeping gene (Gly-tRNA-ligase)detected by qRT-PCR of RDT^+^ subclinical children at different times during the wet (Oct n=14) and the dry season (Jan n=17, Mar n=17, May n=18), and during clinical malaria episodes (MAL n=19). **C** Spearman correlation between Lyso-PC 18:0 and the relative expression of markers of female gametocytes (*pfs25*, left), male gametocyte (*pfmget*, centre), and parasite sexual commitment (*ap2g*, right) . Lipid data and rel expr values compared by Dunn’s Kruskal-Wallis test with Bonferroni multiple comparison correction. Paired samples of the same individuals are connected on plots.

We then sought to relate these data with gene expression of gametocyte-related genes at different timepoints along the year, we used qRT-PCR of female, and male gametocyte markers relative to a housekeeping gene. Our results confirmed that the relative expression of female and male gametocytes markers, although variable, was significantly higher at any timepoint of asymptomatic infection (Oct n =14, Jan n=17, Mar n=17, and May n=18) than in clinical cases of malaria (MAL, n=19) (Fig. 2B). In contrast, no differences were found between the timepoints of asymptomatic infection (Oct, Jan, Mar and May) (Fig. 2B). The expression of *ap2g* was not significantly different across seasons or clinical presentations (Fig. 2B). In addition, no significant correlations were detected between the concentration of LysoPC 18:0 and the expression of *pfs25* (Spearman’s cor 0.286, p-value=0.1474), *pfmget* (Spearman’s cor 0.359, p-value=0.0934) or *ap2G* (Spearman’s cor -0.082, p-value=0.7372) (Fig. 2C), though we did observe a trend of lower LysoPC 18:0 concentrations in clinical malaria cases, that also showed reduced *pfs25* and *pfmget* expression.

### Higher gametocytes proportion in asymptomatic vs clinical infections is independent of seasonality

We then investigated how density and proportion of gametocytes differed between clinical malaria and asymptomatic infections in settings with different seasonality patterns. While Mali experiences contrasting seasons with 5-6 months interruption of transmission every year^10^, Uganda has uninterrupted year-round transmission peaking at times of strong rain^37^. To quantify parasites and female gametocytes in Malian samples as well as proportion of gametocytes out of all iRBCs, we prepared standard curves of asexual parasites and sorted sex-specific fluorescent gametocytes^38^ (Supplementary Fig. 2). Then, we quantified expression of *glycine-tRNA-ligase* and *pfs25* in 19 clinical malaria cases and 50 paired samples (Jan n=25, May n=25) collected from asymptomatic infected individuals at the beginning and end of the dry season, and determined parasitaemia and gametocytemia based on these curves.

To investigate parasite and gametocyte carriage in a non-seasonal setting in Uganda, we obtained total parasite and female gametocyte densities previously published^37^ and retrieved from ClinEpiDB^39^. This analysis included qRT-PCR quantifications of *ccp4* (PF3D7_0903800) and the multilocus *varATS* of 150 individuals with asymptomatic parasitaemias persisting for over one month (n=122) and age-matched individuals with clinical malaria (n=38). In Uganda, we detected 93% of gametocyte prevalence in asymptomatic infections (n=114) and 34% in clinical cases (n=13). In Mali, the prevalence of gametocytes was 69% in the beginning of the dry season in Jan (n=18), and 76% at the end of the dry season in May (n=19) with 11 individuals being positive at both timepoints. Prevalence of gametocytes in clinical cases presented was 89% (n=17). We found significantly higher asexual parasite levels among gametocyte positive samples in clinical malaria cases compared to asymptomatic infections in both settings (Fig. 3A). In addition, a significantly higher density of gametocytes in clinical compared to asymptomatic infections in the Uganda setting was observed, while the difference in Mali was not statistically significant (Fig. 3B). Furthermore, we observed a lower density but higher prevalence of gametocytes (89%) in Malian clinical cases compared to Ugandan clinical samples (34%), suggestion differences in experimental procedure between the two studies might account for the discrepancy in gametocytaemia, with low gametocytaemia in clinical samples remaining undetected in malaria cases of the Ugandan cohort (Fig. 3B). Next, we aimed to determine the proportion of gametocytes of the total parasite population in both settings. Our data showed that in both Mali and Uganda, gametocytes constitute a larger proportion of the total parasite population in asymptomatic infections than in clinical cases of malaria (Fig. 3C), in line with the described increased relative expression of gametocyte-marker genes (Fig. 1 and 2). On the other hand, transcription of *ap2g* was comparable in the Malian clinical and asymptomatic samples (Fig. 3D). Also in the Malian setting, paired samples from the beginning and end of the dry season revealed a significant decrease of parasite load from January to May (Fig. 3E), while no significant difference in gametocyte levels (Fig. 3F), or commitment rate measured by *ap2G* expression were detected (Fig. 3G), supporting unaltered sexual commitment or gametocyte development throughout the dry season.

**Figure 3.**
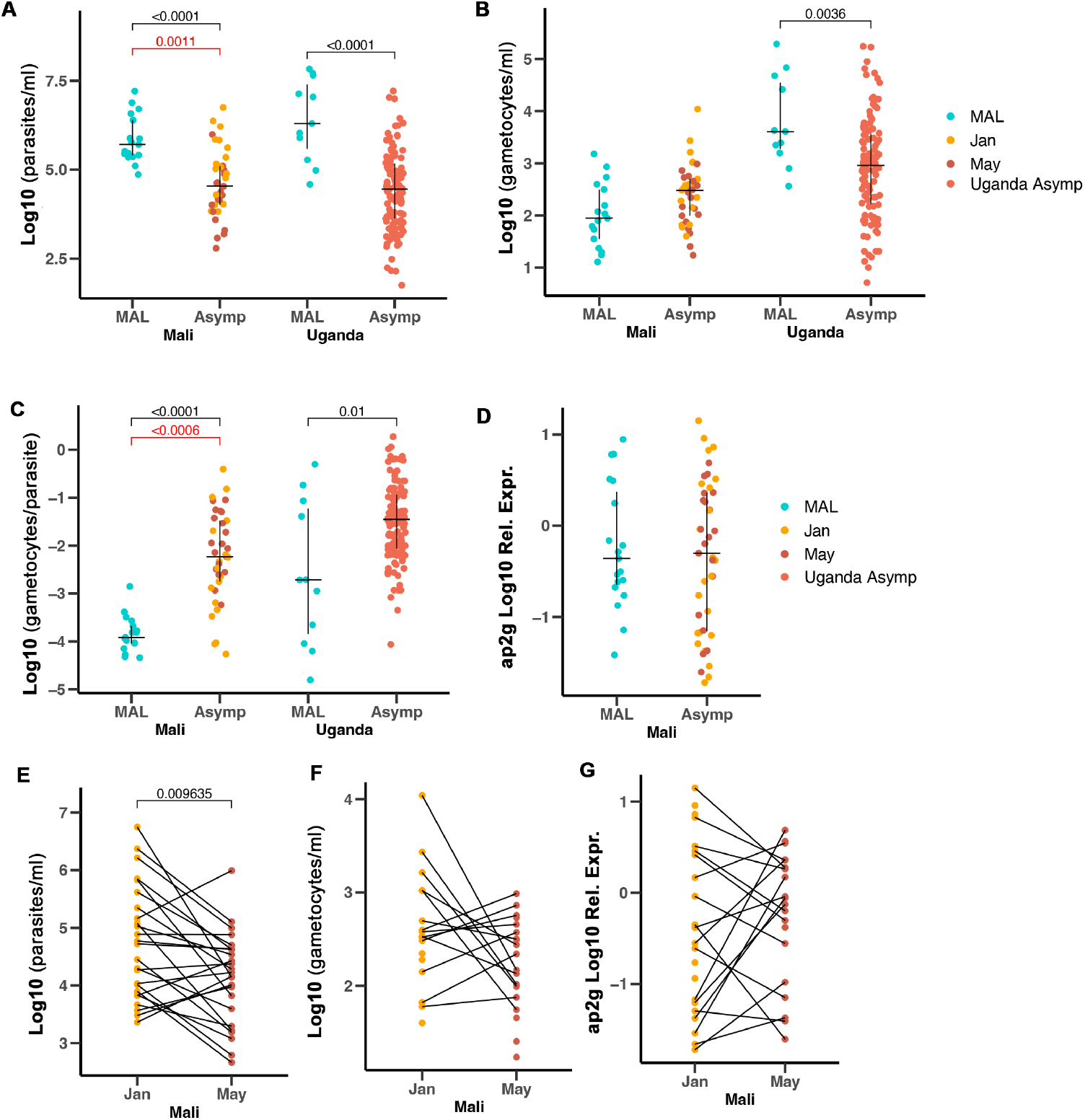
Proportion of gametocytes is higher in asymptomatic infections than in clinical cases. **A** Parasite density detected by qRT-PCR of *Glycine-tRNA-ligase*, using a standard curve of Pf2004 parasites and sorted female gametocytes, in gametocyte positive symptomatic (n=13, MAL) and asymptomatic infections (n=113) in Uganda; and clinical malaria cases (MAL n=17), and asymptomatic infections (Jan n=18, May n=19) from the beginning and end of the dry season in Mali. **B** Gametocyte density in the same samples quantified by qRT-PCR of *pfs25* relative to a standard curve of sorted female gametocytes. Samples from Uganda and Mali were compared separately. p-values for the MAL-Jan comparison and Jan prevalence in Mali samples are shown in red, MAL-May comparison shown in black. Values are plotted with median and IQR Dunn’s Kruskall-Wallis test with Bonferroni multiple comparison correction. **C** Ratio of gametocytes to asexual parasite load. Values are plotted with median and IQR, Dunn’s Kruskall-Wallis test, samples from Uganda and Mali were compared separately. P-values for the MAL-Jan comparison are shown in red, MAL-May comparison in black **C** Relative expression of *ap2g* in malaria cases (n=19) and asymptomatic dry season (Jan n= 20, May n=18) from the same sample set, Dunn’s Kruskall-Wallis test. **D** Parasite density, quantified by Glycine-tRNA qRT-PCR relative to a standard curve for 25 paired samples from the beginning (Jan) to end of the dry season (May). Wilcoxon matched-pairs signed rank test. **E** Gametocyte density quantified by *pfs25* qRT-PCR relative to a standard curve in samples from the beginning (Jan, n=18) to end of the dry season (May, n=19). Wilcoxon matched-pairs signed rank test performed on 11 paired samples with detectable gametocytes in both timepoints. **F** *ap2g* expression by qRT-PCR of samples from the beginning (Jan, n=20) to end of the dry season (May, n=18). Wilcoxon matched-pairs signed rank test performed on 18 paired samples with detectable *ap2G* expression at both timepoints.

### Progressing infection leads to apparent differences in gametocytogenesis

We observed higher proportions of gametocytes in asymptomatic vs clinical malaria, without concomitant differences in the expression of markers of sexual commitment and early gametocytes. As infection progression over the course of the dry season, we observe a decrease in parasitaemia, though no apparent change in the proportion of gametocytes. To explain this, we devised a mathematical model considering three different parasite populations - asexual parasites (P), sexually-committed parasites (PG) and gametocytes (G), all of which circulate or are cytoadherent^31^ (Fig. 4A). To improve the fit of an earlier model by Cao et al.^31^ to our data, we extended the model by incorporating the assumption of increased clearance of longer-circulating iRBCs in persisting parasitaemias compared to recently transmitted parasites as reported previously^11^. This distinguished two phases of infection, in which the growth of parasites varied due to immune response or the parasite’s adhesion properties, in agreement with reported data^37^. In our model, parasitaemia increased rapidly at early timepoints after parasite egress from the liver (day 0), up to day 14 when, in our model, infections transition to the chronic phase. Faster clearance in the chronic phase due to immunity or splenic removal of parasites was implemented as an increase in the asexual parasite removal at the transition from acute to chronic infection on day 14. We then followed chronic infections up to day 180, mirroring the 6-month dry season length in Mali. We fit the model to the symptomatic and asymptomatic parasitaemia and gametocytaemia levels of the Malian cohort reported in Figure 3, assuming that clinical cases occur shortly after transmission of parasites on day 14 of blood stage as evidenced by epidemiological studies associating disease with recent transmission^12,40,41^. Asymptomatic infections at the beginning of the dry season, i.e. January, were used to describe early asymptomatic timepoints (day 28), and the end of the dry season in May was used as a late asymptomatic timepoint (day 180) (Fig. 4C and D). Using an iterative Bayesian fitting-approach to fit the model to the symptomatic and asymptomatic parasitaemia and gametocytaemia levels of the Malian cohort, we quantitatively assessed biological parameters including sexual commitment rate, gametocyte circulation time, and gametocyte sequestration time, confirming previously determined parameter ranges^31,42^ (Supplementary Table 4 and Fig. 4E). Our adapted model provided a dynamic depiction of parasitaemia levels, as well as gametocyte densities and proportions, as seen in the Malian data. In line with our data and previous literature ^43^, parasitaemia in chronic infection decreased following a log-linear pattern until day 180 (Fig. 4B).

**Figure 4.**
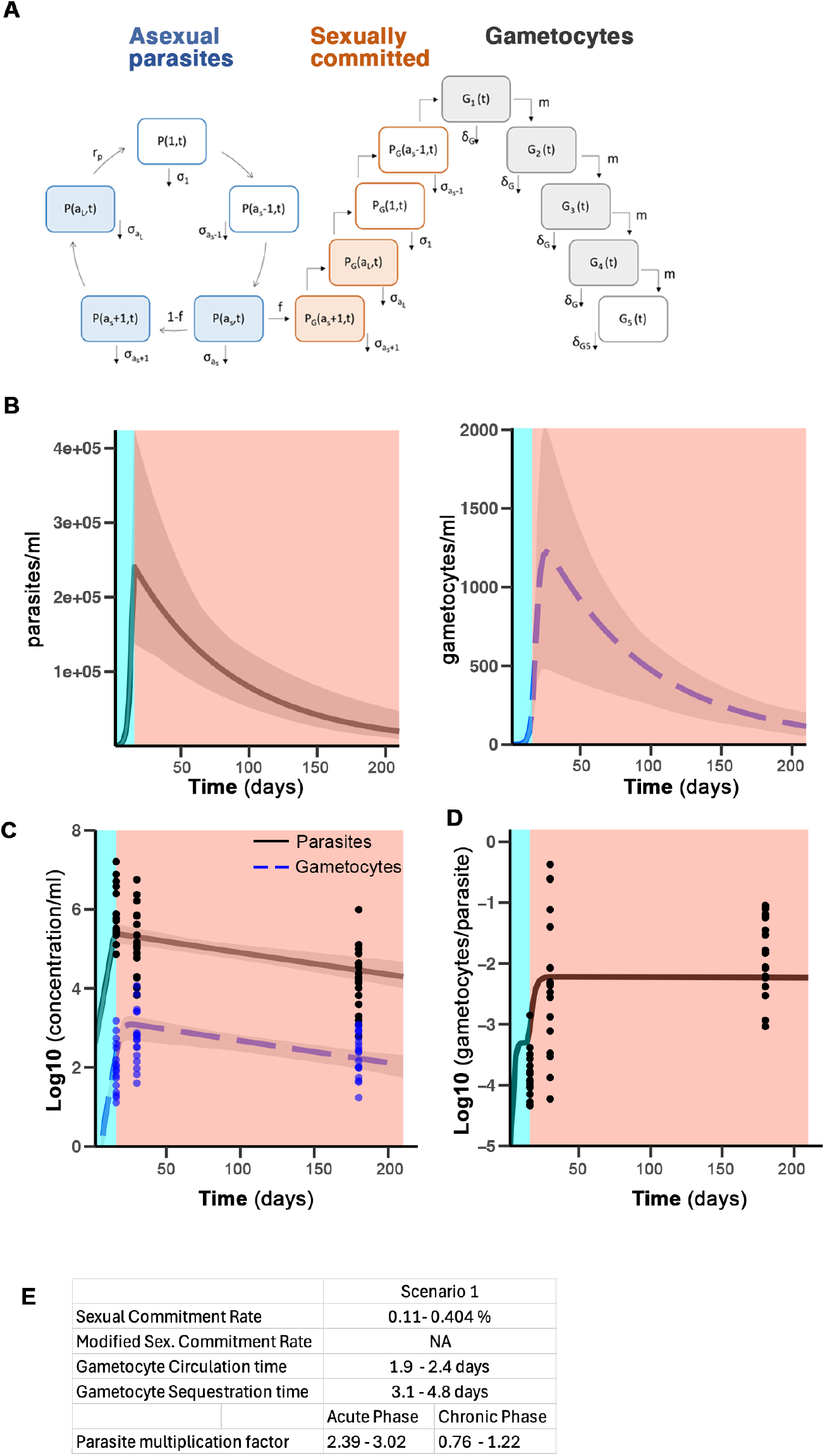
**A** Sketch of the mathematical model to describe parasite dynamics including individual compartments, transitions and death rates: Asexual parasites (P) replicate in a 48h cycle, being cleared at an age dependent death rate (σa). A fraction of asexual parasites (f) transitions to the sexually committed parasite compartment (PG), goes through a round of replication and develops into gametocytes (G). Gametocytes mature over 5 distinct stages. Asexual parasites (blue), sexually committed parasites (orange) and gametocytes (grey). **B** Model prediction of development of parasitemia (left) and gametocytemia (right) with infection progression. Parasitemia and gametocytemia were averaged over one replication cycle (48h). Solid and dashed lines show average of model predictions based on posterior parameter distribution, with grey shaded areas corresponding to the range of model predictions. **C** Parasitemia and gametocytemia. Lines indicate model prediction, points refer to the actual measurements of gametocytemia for MAL (d14), Jan (d28) and May (d180). **D** Ratio of gametocytes/parasites as predicted by the model (lines) and individual measurements for clinical malaria cases (d14), Jan (d28) and May (d180). **E** Parameter estimates in distinct phases of *P. falciparum* infection shown as 89% credibility interval of the posterior distributions. The sexual commitment rate defined by the fraction of parasites that are sexually committed, *f*, is given in %.

Maintaining the fraction of parasites entering sexual commitment, *f*, constant throughout acute and chronic infection phases, our model predicted a lower proportion of gametocytes in the acute phase (Fig. 4D), when parasitaemia increased rapidly. Due to the maturation time of gametocytes outside of circulation, gametocytemia lags behind parasitaemia, as gametocyte levels are reflective of parasite density at the time when sexual commitment occurred. This trend is reversed during the chronic phase of infection due to the negative growth of asexual parasites imposed by increased removal of iRBCs. After a transition phase when the proportion of gametocytes in the population increased, the gametocyte proportion reached a higher equilibrium (Fig. 4D). Therefore, a model with a fixed commitment rate seemed capable of reproducing the patterns of parasitaemia and gametocytaemia as we observed in the Malian data.

To further test how alternative assumptions would affect data explainability, we investigated how a dynamic commitment rate would influence parasite and gametocyte dynamics as infection progresses. To this end, we modified the model framework to investigate two additional scenarios, in which the commitment rate changes as infection progresses. Scenario 2 assumes that the sexual commitment changes during the persistent phase of infection, i.e. at day 100 post infection, reflecting potential changes due to seasonal cues (Supplementary Fig. 3A). In contrast, scenario 3 allows for a varying commitment at the time of peak parasitaemia, i.e. between day 7 and 21, mimicking the potential impact of Lyso-PC depletion in the blood stream (Supplementary Fig. 3B). Using the same Bayesian fitting-approach, we fitted the two alternative models to the data and compared their descriptive power (Supplementary Fig. 4A) and inferred parameter ranges (Supplementary Fig. 3C). Our original (scenario 1) and scenarios 2 and 3 were able to explain the observed dynamics with similar posterior distributions for all model parameters other than the commitment rates (Supplementary Table 4, Supplementary Fig. 5A).

Commitment rates in scenario 1, 2 and 3 fell in a similar range. Although in scenario 2 and 3 the range of estimated commitment rates in the persistent phase of infection were wider and also showed tendencies towards higher estimates if algorithms were run for longer, they still overlapped with the constant commitment rate assumed in scenario 1 with differences of the commitment rates between different scenarios remaining minor in absolute terms. Thus, the data do not contain enough support for a scenario that assumes a changing commitment rate, as a fixed commitment rate is equally likely to provide a dynamic depiction of the observed gametocyte densities and proportions and is generally preferred by the model selection criteria (Supplemental Figure 5).

## Discussion

The absence of mosquitoes during dry months and the limited lifespan of *P. falciparum* gametocytes^8,9^ renders the production of gametocytes useless during the dry season. However, gametocytes must be available to resume malaria transmission when mosquitos return each wet season. Whether *P. falciparum* maintains a constant rate of producing sexual forms or adjusts gametocytogenesis to optimize transmission and fitness is unclear. In this study, we used samples from Malians exposed to alternating six-month dry and wet seasons, as well as samples from clinical malaria and persisting *P. falciparum* infections from Uganda, where transmission occurs year-round. Our findings demonstrate that clinical malaria cases show higher parasite and similar or higher stage V gametocyte densities compared to asymptomatic infections. However, the proportion of gametocytes is increased in asymptomatic persisting infections. Additionally, we observed that transcription of genes characterizing sexually-committed parasites and stage I gametocytes^44,45^ were unaffected by season or symptoms. This clearly contrasts with genes of late-gametocytes, which were enriched among upregulated transcripts in persisting infections at the end of the dry season. Together these observations suggest that gametocytogenesis remains constant as infections progress from recent transmission to chronic carriage, and it is the difference in asexual survival/replication rates that promotes different gametocyte levels.

It has been suggested that *P. falciparum*’s investment in gametocyte production adjusts to host nutrient availability^46^, drug pressure^17^, transmission intensity^47^, host inflammatory state^48^, and seasonality^49,50^, but results are inconsistent and potentially affected by the dynamics of asexual parasite growth. In Sudan, it was observed that gametocyte density increased prior to the highest transmission peak^51^, suggesting higher gametocyte investment. In Burkina Faso, gametocyte densities were higher at the peak and end of the transmission season than during the dry season^52^, while the efficiency of mosquito infection through membrane feeding was greater early in the transmission season than at the peak of the wet or the dry seasons^53^. Nevertheless, these studies did not monitor the same parasite population longitudinally from the dry into the wet season. Also, in Burkina Faso, transmission was more effective in persistent than in incident infections, but data collection ended after day 35, and all collections were done during the wet season^18^.

In vitro studies have shown that depleting Lyso-PC or choline induce *ap2G* expression and increased gametocyte production^25,54^, but associations in vivo are less consistent. In Rwanda, low Lyso-PC levels were associated with higher parasite densities^55^, while in Ghana, although correlations with parasitaemia were not reported, Lyso-PC levels inconsistently associated with gametocyte-committed rings converting into gametocytes in vitro^45^. Our data, similar to Orikiiriza et al.^55^, demonstrate an association between lower Lyso-PC levels and increased parasitaemia and malaria symptoms. However, this association was lost at low parasitaemias, and Lyso-PC levels were unaffected by asymptomatic carriage compared to non-infected individuals. In line with previous reports of parasitaemia affecting lipid levels by its consumption for parasite growth^25^. In vitro, drugs also affect sexual conversion; sub-curative doses of dihydroartemisinin (DHA) stimulate *ap2g* expression and sexual conversion of trophozoite-, but not ring-stages. But this increase could not synergize with choline depletion, suggesting that the effects of DHA on lipids, might be the same as choline depletion^54,56^. In vivo, it has been shown that after artemisinin treatment, *ap2g* expression increases in many, but not all patients, and data varied significantly between cohorts. These findings highlight the complex regulation of sexual conversion which is influenced by patient, parasite characteristics^57^, and resistance status^58^. Our RNA-seq and RT-qPCR data reveal a constant expression of *ap2g* and unaltered expression of transcripts associated with committed schizonts and early-gametocytes, suggesting that reduced Lyso-PC observed during uncomplicated clinical malaria is unable to affect *P. falciparum* sexual commitment. Accordingly, in vitro supplementation with plasma from clinical malaria cases or end of the dry season infected individuals did not differentially affect the parasite’s sexual conversion rate. This supports that Lyso-PC differences introduced by *P. falciparum* infection are not sufficient to trigger the mechanisms demonstrated in vitro^25,54^. Nevertheless, it is possible that in specific niches Lyso-PC levels are sufficiently low to trigger the sexual commitment of parasites, although this requires the presence of iRBCs in such niches to respond to the stimulus.

Previous studies on gametocytogenesis did not consider the dynamic changes within the asexual compartment during persistent infections. Greischar et al. demonstrated the difficulty of quantifying transmission investment by comparing existing quantification methods on simulated data with known transmission investment^59^, while Prajapati and colleagues relied on in vitro culture of sexually committed circulating parasites to define samples with varying commitment rates^60^. Also, our study has limitations; the quantification of parasite and gametocyte densities may be affected by primer efficiencies, which may impact comparisons across different transcript measurements, although our relative analysis to a standard curve minimizes such effects. Although we lack longitudinal data and make assumptions about the timing of clinical and asymptomatic infections during infections progression, and while our model only represents the dynamics of infections with a single parasite strain, our model can explain the observed patterns by assuming transitions from a faster to a slower asexual growth rate^61^. Despite these limitations, we show that the proportion of sexual stages is inversely related in recent transmissions versus chronic infections, without alteration in the expression of regulators of sexual commitment while maintaining a constant rate of sexual commitment. Taking into account that persisting infections beyond the acute immune response exhibit prolonged circulation and increased splenic clearance^11^, allowed us to complement the model by Cao et al.^31^ with an increased asexual parasite death rate following the immuno-responsive period based on reports of prolonged circulation and increased splenic clearance^11^. This promotes a progressive increase in the representation of sexual stages in later phases of infection, which recent data showing peak of gametocyte density preceding by several weeks the peak of gametocyte proportions, also support^22^. The low gametocyte proportions early in infection are due to the lag of ∼8-12 days of gametocyte development in the bone marrow, while asexual replication efficiently increases in the absence of adaptive immunity. Data of controlled infections has indeed shown that parasitaemia is predictive of gametocytemia with a ∼14-day delay^29^. Later, immunity prevents fast asexual growth and parasites that persist further are more effectively removed in the spleen leading to sustained low parasitaemias. The ∼5-12-day lag has the opposite effect later in infection when asexual parasitaemia decreases and the proportion of gametocytes reflects the parasitaemia of ∼5-12 days earlier; all despite maintaining the same sexual commitment rate.

In conclusion, our data is consistent with a model in which individuals not undergoing major lipid deprivation, present a constant rate of *P. falciparum* sexual commitment as infections progress from recently transmitted to persistent over several months.

## Methods

### Gametocyte gene expression analysis

We used RNAseq data of leucocyte-depleted blood pellets from 12 children with asymptomatic malaria at the end of the dry season and an age- and sex-matched cohort presenting with their first malaria case in the ensuing transmission season reported in^11^. Originally 1456 gametocyte specific genes were reported^32^, and we counted 1258 after excluding repetitions and converting nomenclature to its most recent version (Supplementary Table 1). Out of these 1258 genes, we detected normalized reads of 551 gametocyte-specific genes in the RNAseq data set^11^ to perform a principal component analysis.

### Analysis of early and late gametocyte markers in RNA-seq data

We compiled a list of early and late gametocyte specific genes based on three RNA-seq studies comparing distinct parasite stages, including data from Poran et al. 2017^33^ obtained after request from the corresponding author (compared day 2 versus day 9 of gametocytogenesis by single cell RNAseq); data from Lopez-Barragan et al. 2011^34^ (compared day 4 versus day 15 of gametocytogenesis by bulk RNAseq) and data from Jeninga et al.^35^ (compared day 4 of gametocytogenesis versus day 10 of FACS sorted male and female gametocytes by bulk RNAseq). Poran et al. selected early and late gametocyte genes as genes showing a 20-fold higher expression in early or late gametocytes over the respective other group. Late gametocyte genes were also required to show a 20-fold higher expression in late gametocytes compared to ring stage parasites, this was not the case for early gametocyte genes. In Lopez-Barrgagan’s data, early gametocyte genes were defined as genes having a 10-fold expression increase in early gametocytes compared to late gametocytes. Genes showing a 20-fold higher expression in late stages were designated as late gametocyte genes. Expression differences to asexual stages were not considered. We revisited Jeninga et al. RNA-Seq data for ring stage parasites, schizont stage parasites, day 4 and 10 male and female gametocytes from GSE180985 and evaluated differential expression of sense transcripts. A countmatrix was generated using Cufflinks (REF: doi:10.1038/nbt.1621). The matrix was analysed by filtering for genes with at least 3-fold differential expression, and a cutoff of minimum 10 FPKM per gene was applied. Intersecting the full list of all three publications describing early and late gametocyte genes with genes transcribed in circulating parasites stages, defined by RNAseq data of Andrade et al., generated a final list of 109 early and 172 late gametocyte genes to evaluate expression differences of these genes between dry and rainy seasons. Intersecting the early and late gametocyte list and all gametocyte genes reported by Painter et al., we obtained 551 gametocyte genes that were detected in the wet vs dry season RNAseq dataset^11^. Of these 343 were among differentially expressed determined using Deseq2 ^62^ (padj < 0.05, LFC >1). For GSEA, differentially expressed genes between MAL and May were ranked by Log2 FC and analysed with the fgsea ^63^ algorithm implemented in the clusterprofiler ^64^ R-package.

### Study subjects and ethical approval

Approval by the Ethics Committee of Heidelberg University Hospital, the Faculty of Medicine, Pharmacy and Odontostomatology (FMPOS) at the University of Bamako, and the National Institute of Allergy and Infectious Diseases of the National Institutes of Health Institutional Review Board was obtained to conduct the study registered at ClinicalTrials.gov with the identifier NCT01322581. Clinical data and samples were obtained from 2012 to 2019 in a cohort study previously described^65^, conducted in Kalifabougou, Mali, where malaria transmission occurs between June and December, and antimalarial drugs are provided by a single clinic and pharmacy. All participants and the parents/guardians of included children provided written informed consent. Exclusion criteria at enrolment included haemoglobin <7 g/dL, axillary temperature ≥37.5°C, acute systemic illness, or use of antimalarial or immunosuppressive medications in the 30 days preceding enrolment. Clinical malaria cases were prospectively determined by passive surveillance and defined by axillary temperature ≥37.5°C, ≥2500 asexual parasites/μL of blood, and no other evident cause of fever. Malaria episodes were treated with a 3-day standard course of artemether/lumefantrine according to Malian national guidelines. Cross-sectional clinical visits and blood draws were performed late in the transmission season (October), and at the beginning (January), mid (March) and end (May) of the dry season.

### Sample collection

Venous blood (4 mL if donor age was below 4 years, and 8 mL if above) was drawn by venepuncture on wet and dry season cross-sectional visits, and at first malaria episode of the transmission season. Samples are referred as MAL for malaria cases or by time point of collection. Blood was collected in sodium citrate-containing cell preparation tubes (Vacutainer CPT Tubes, BD) and transported to the laboratory where RBC pellet and plasma were separated and stored at −80°C within three hours of blood draw. Plasma used for lipidomic analysis and induction of gametocytogenesis was separated by centrifugation and immediately frozen in liquid N2.

### Parasite culture and preparation of a standard curve for gametocyte quantification

Pf2004 parasites were cultured at 37° C in 5% O2, 5% CO2 and 90% N2, at 5% haematocrit in RPMI-1640 medium (with L-glutamine and HEPES (Gibco)). Parasite culture medium was supplemented with 0.24% Sodium Bicarbonate (Gibco), 100μM Hypoxanthine (cc pro), 25 µg/mL gentamycin (Gibco)), and 0.25% Albumax II (Gibco). Medium was changed daily, parasitaemia assessed by Giemsa-stained blood smear and the culture split according to experimental needs.

*PfDyn*GFP/*PfP47*mCherry parasite lines^66^ were cultured in presence of 15 μg/ml of Blasticidin-S-HCL (Invitrogen) and 2 μM Pyrimethamine (Sigma Aldrich) as described for non-integrating plasmids and P52 integrating plasmid. Parasites were grown to >4% parasitaemia when 50 mM N-Acetylglucosamine (Sigma Aldrich) was added to the culture to eliminate asexual stages. After 4 consecutive days of N-Acetylglucosamine treatment, culture in complete RPMI was resumed. Gametocyte development was monitored daily by smear, and on day 5 after start of gametocyte induction magnetic enrichment was performed with MACS columns (Miltenyi Biotec). Gametocytes were sorted on a BD FACSAria III on day 8, keeping cells at 4° C in suspended activation buffer, (gating strategy in Supplementary Figure 2A). RNA of a defined numbers of sorted male and female gametocytes was extracted and reverse transcribed, a standard curve prepared by serial dilution and used for RT-qPCR as described below.

### RNA extraction and quantification of parasite transcripts

RNA was isolated from 200 µl RBC pellet by phenol-chloroform extraction using TRIzol (Ambion), followed by DNAse treatment using the DNAse I Amplification Grade Enzyme (Invitrogen). RNA was quantified by absorption measurements at 230, 260 and 280 nm on a Citation3 Imaging plate reader, or using the Qubit RNA High Sensitivity kit (Thermo Scientific) followed by cDNA synthesis using the Superscript IV Enzyme (Invitrogen) according to manufacturer instructions. A Taqman RT-qPCR multiplex assay (Applied Biosystems) was performed on a qTower (Analytic Jena) with primers to quantify the female gametocyte marker ookinete surface protein *pfs25* (PF3D7_1031000), the male gametocyte transcript *pfmget* (Pf3D7_1469900), the master regulator of sexual commitment *ap2g* (PF3D7_1222600) and a parasite housekeeping gene (*glycine-tRNA-ligase*, PF3D7_1420400). We quantified primer efficiency and specificity with an NF54 parasite line expressing DyneinGFP and carrying an episomal P47mCherry plasmid (DynGFP/P47mCherry) was used for sorting of male and female gametocytes through their respective specific mCherry and/or GFP expression ^38^.

(Supplementary Figure 2a)

We generated standard curves of female or male gametocytes and asexual parasites by serial dilution of cDNA of 1*10^6^ FACS-counted male and female gametocytes and asexual parasites in 10-fold dilution steps down to 10 parasites/gametocytes per ml. Using the standard curve, amplification efficiency (E) was estimated using the equation 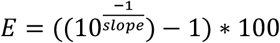 (Supplementary Figure 2B). *pfs25* and *pfmget* expression was absent in asexual parasite and high in sorted gametocytes was high and decreased with increasing dilution (Supplementary Figure 2b). *pfmget* was approximately 10-fold higher expressed in male gametocytes (double+) compared to females (mCherry+), consistent with 5-7% sorting impurity. (Supplementary Figure 2b) Log-linear regression fit for the independent populations revealed a good fit with R^2^-values of 0.94-0.98 for *ap2-g*, 0.99 – 0.998 for *glycine-tRNA ligase*, 0.97-0.99 for *pfmget* and 0.997 -0.999 for *pfs25*. Primer efficiencies for the 4 genes ranged from 97% (*pfs25*) to 115% *ap2g*) and were corrected for in qPCR quantification. The combination of asexual and sexual standard curves was used for qPCR quantification of total parasitaemias and female gametocytemia throughout the season. To account for fluorescence leakage between channels a compensation matrix was calculated based on single colour fluorescence measurements by an application specialist at Analytic Jena. *Gexp5* (PF3D7_0936600) expression levels were quantified in a Taqman RT-qPCR duplex assay together with *glycine-tRNA ligase* as a housekeeping gene. All primers and probes sequences are listed in Supplementary Table 6. Relative quantification was performed by ΔΔCt method using *glycine-tRNA ligase* as housekeeping gene and May samples a calibrator group. For all genes except *gexp5*, primer Pfaffl correction was used in the quantification to account for different primer efficiencies ^67^.

### Phospholipid analysis

May and MAL plasma samples (shown in Figure 1) were analysed by shotgun lipidomics at Lipotype as described in^68^. Briefly, automated samples extraction was followed by direct sample infusion and high-resolution Orbitrap mass spectrometry including lipid class-specific internal standards to assure absolute quantification of LysoPC and LysoPE, using LipotypeXplorer for identification of lipids in the mass spectra. Lipid species detected less than half of samples were excluded, concentrations were normalized to total lipid amount in each sample. Oct, Mar, May and MAL samples (shown in Figure 2 A,C) were analysed by Liquid chromatography-mass spectrometry (LC-MS). Samples were extracted, processed, and analysed at the Lipidomics Unit at the Institute of Physiological Chemistry in Mainz, using the liquid-liquid extraction protocol as described^69^. All values were normalized to the analysed plasma volume.

### In vitro induction of *P. falciparum* sexual commitment and gametocytogenesis

In vitro induction of *P. falciparum* sexual commitment was performed as described by (Brancucci et al., 2015). Pf2004 parasites were synchronized to a 4 h window by repeated treatment with 5% sorbitol and MACS purification. Parasites at 26 ± 2 h post invasion (hpi) were plated on 48-well plates in 500 µl culture volume at 0.8-1% parasitaemia and 2.5% haematocrit and cultured in minimum fatty acid medium (mFA, RPMI1640 with 100 µM hypoxanthine, 25 µg/ml gentamycin, 0.39% fatty-acid-free BSA, 30 µM palmitic acid and 30 µM oleic acid) supplemented with 25% plasma from asymptomatic children at the end of the dry season (PlMay), or 25% plasma from malaria cases during the rainy season (PlMAL), in addition to the conditions used by Brancucci et al. (mFA alone, and mFA supplemented with 25% German plasma (PlDE)). After 24 h of treatment, the medium was changed to culture medium and subsequently changed daily. At 48 h after beginning of treatment 30 IU/ml heparin were added to block re-invasion of parasites and eliminate asexual forms. At 0, 48 and 120 h post treatment RBC pellets were collected for parasitaemia and gametocytaemia determination by flow cytometry (50 μl of culture stained with 5x SYBR Green II (Invitrogen) and 7,5 μM MitoTracker (Applied Biosystems) for 30 min at 37 °C and acquired with CountBright Absolute Counting Beads (Thermo Ficher) on a FACS Canto II or an LSR II and analysed using FlowJo software 10.2 or higher versions (Tree Star)). Additionally, Giemsa-stained blood smears were prepared at 48 h and 120 h post-treatment and blindly analysed by microscopy, counting RBCs, parasites, and gametocytes in 18 fields of view. Multiplication rate of parasites was calculated as 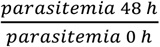, and conversion rate as 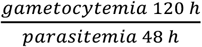.

### Quantification of parasites in samples of Ugandan asymptomatic carriers by RT-qPCR

We compared the Malian data from dry and wet season with data from 150 Ugandan individuals enrolled in a 2-year longitudinal study including monthly standardized clinical evaluation and blood collection^37^ retrieved from ClinEpiDB (https://clinepidb.org/ce/app/record/dataset/DS_51b40fe2e2122). Asymptomatic *P. falciparum* positive individuals (20.4 years ± 17.2, median: 13.3 years) were included based on the following criteria: i) ≥ two PCR+ time points, ii) ≤ 3 negative time points in between positive, iii) same clone (determined by deep amplicon sequencing of atypical membrane antigen 1 (*ama1*) as described in^70^ present at all time points of one asymptomatic infection. Additionally, we used samples from an age-matched group of 38 clinical malaria cases (tympanic temperature > 38°C or history of fever in the previous 24 hours and *P. falciparum* positivity by microscopy). *P. falciparum* parasite and gametocyte density in blood samples were measured by qPCR of multi copy conserved *var* gene acidic terminal sequence^71^ and *ccp4* (PF3D7_0903800), respectively. Average levels of parasitaemia and gametocytaemia during each asymptomatic infection, were estimated as the area under curve over the course of infection divided by length of infection. For symptomatic infections, only the quantification before treatment on day of diagnosis was considered.

### Mathematical model

The developed mathematical model extends a model used previously to describe asexual parasite and gametocytes dynamics ^31^. The model considers 3 different main parasite populations, i.e., distinguishing between asexual parasites (*P*), sexually committed parasites (*P*_*G*_) and gametocytes (*G*) (Fig 4E). Asexual parasites develop and replicate within a red blood cell over a cycle of length *a*_*L*_ =48h after which mature schizonts burst and release a number of *r*_*P*_ =18-32 merozoites that are able to invade new red blood cells. During this replication cycle, parasites are assumed to be removed through splenic clearance and active immune response. In extension to the previous model, we assume that this removal of asexual and committed parasites follows an age dependent death rate, *σ*_*a*_ that differs in distinct phases of acute and chronic *P. falciparum* infection. The following iterative equation describes asexual parasite growth at discrete time steps for parasite age, *a*, and time, *t*, increments of one hour.

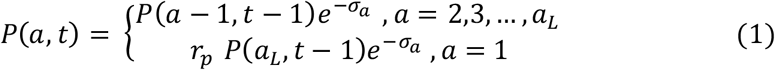

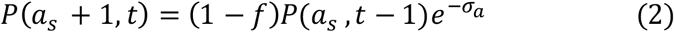

The rate of parasite removal, *σ*_*a*_, is assumed to increase with parasite age within one replicative cycle and is described by a sigmoidal curve given by:

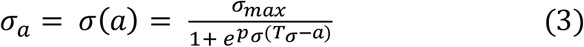

The parameter *σ*_*max*_ defines the maximal rate of parasite removal, *T*_*σ*_, the age at which 50% of parasites are removed, and *p*_*σ*_ a scaling parameter.

At parasite trophozoite stage of *a*_*s*_ = 20hpi parasites adhere to the endothelium when at the same time a fraction, *f*, of parasites enters sexual commitment. Committed parasites continue the rest of the replication cycle producing only committed parasites that enter the first gametocyte stage at the sequestration age, *a*_*s*_, of the following replication cycle

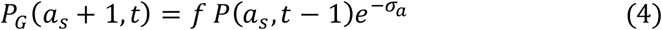

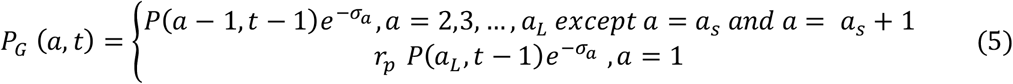

Gametocyte development through five stages, of which only stage 5 is found in circulation, is modelled by:

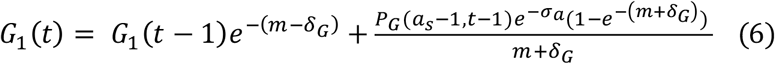

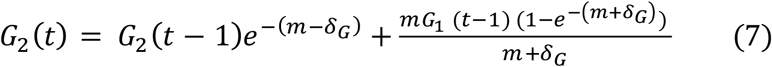

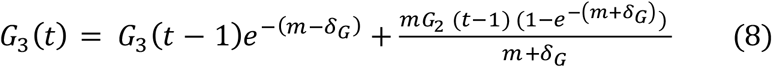

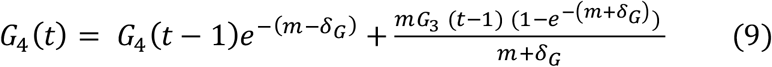

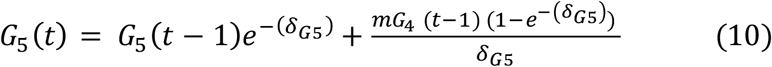

Immature gametocytes, that are sequestered, mature with a maturation rate, *m*, and are lost by death rate *δ*_*G*_, while mature circulating gametocytes are removed by a splenic clearance rate *δ*_*G*5_. As pointed out above the whole model is simulated by discrete time-steps of one-hour increments.

Our model assumes two stages of *P. falciparum* infection, accounting for an acute and a chronic asymptomatic phase. The start of the infection is marked by the beginning of blood stage infection, for which we assume an initial parasite load, *IPL*, with parasite age normally distributed with mean, *μ*_IPL_ and standard deviation *σ*_IPL_. During the acute phase of infection parasitaemias increase rapidly leading to clinical symptoms within the first 2 weeks of blood stage infection^72^. At 14 days after the start of blood stage infection, the maximal parasite removal rate is assumed to increase to *σ*_*max*2_ to account for the onset of an effective adaptive immune response and, thus, an increase in age-dependent parasite loss. This timepoint indicates the beginning of the chronic phase of infection.

The model described in Eqs (1-10) was applied to the data on symptomatic and asymptomatic log-transformed parasitaemias and gametocytemias of the Malian cohort assuming a fixed parasite life-cycle period of *a_L_*=48h, a replication factor of 25, a sequestration age of *a_s_*= 20hpi, as well as *T_σ_*=32h and *P_σ_*=0.7 characterizing age-dependent parasite removal during the acute infection phase. For the initial parasite load, *IPL*, we assume a fixed Gaussian parasite age distribution with parasite mean age *m_IPL_*=7h and standard deviation *σ_IPL_*=5.5. All remaining parameters were fitted to both symptomatic and asymptomatic time points simultaneously using an approximative Bayesian computing approach (pyABC, Klinger et al. Bioinformatics 2018 ^73^). Prior distributions for the individual parameters spanning biologically realistic ranges (Supplementary Table 2), as well as all fixed parameters (Supplementary Table 3) were selected in accordance with previous studies. pyABC was run with the following stopping parameters: *ε*_*min*_ =0.01, maximal population size *pop*_*max*_ =100, Δ*ε*_*min*_ = 0.05. Given the complexity of the model and the available data, parameter posterior distributions are reported for the 10^th^ pyABC-generation of several independent pyABC fitting runs in order to check robustness of results and avoid local convergence of algorithms (Supplementary Figure 3). Using the Δ*ε*_*min*_ -criterion for stopping did not change the overall conclusions.

To test model sensitivity with regard to the fixed parameter values, we examined the effect of using the high and low boundaries for the reported estimates for each of the constants on posterior parameter distributions. While expected dependencies of the replication rate, *r*_*P*_, and *σ*_T_ with the removal rates *σ*_max_ and *σ*_max2_, could be observed, the overall posterior parameter distributions for the free parameters proved stable to changes in model constants (Supplementary Fig. 4).

The parasite multiplication factor was calculated by *r*_*p*_*e*^−*total clearance*^(1 − *f*) with the term *e*^−*total clearance*^ describing cumulative clearance per replication cycle. Gametocyte sequestration time was calculated by 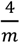 in relation to the 4 sequestered states (stage I to IV) ^74^.

For scenario 2 and scenario 3, the model framework was modified by assuming a change in the fraction of parasites entering sexual commitment, *f*, at different timepoints. In scenario 2, *f* was assumed to change to *f*_2_ at day 100 after infection, while in scenario 3, we assumed a deviating fraction *f*_3_ from day 7 to day 21. All other parameter and model constants were left unchanged, and parameter fits were performed with the same pyABC-settings as before, assuming the same prior distributions for *f*_2_ and *f*_3_ as for the parameter *f*.

For model comparison, the Bayesian Information Criterion (BIC) was calculated by *BIC* = −2LL +*klog*(*n*) with *k* being the number of free parameters and *n* the number of data points. The log likelihood was calculated as the sum of each gametocytemia and parasitemia measurement calculated using a normal distribution as 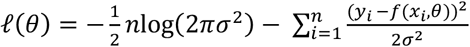where *σ*^2^ is the variance, *y*_*i*_ the corresponding measurement at time point , *x*_*i*_ and *f*(*x*_*i*_, *θ*) the model output. The BIC in scenario 1 with an unchanged commitment rate was slightly lower than in the other two scenarios, due to the lower number of free parameters (Supplementary Fig. 5A). In general, several of the model parameters cannot be independently determined showing correlations between several of the parameter estimates including between the initial parasite load *IPL* and the removal rates *σ*_max_ and *σ*_max2_, respectively, as well as between the two removal rates *σ*_max_ and *σ*_max2_ (Supplementary Fig. 5B). These correlations are expected from the model structure inhibiting their independent inference. Of note, we did not observe a strong correlation between the fraction of committing parasites *f, f*_2_, and *f*_3_ and any of the other parameters (Supplementary Fig. 5B). We assessed robustness of the parameter estimates obtained by pyABC by running the fitting algorithm 100 times for each of the three scenarios and compared parameter estimates in the 10^th^ generation. For all three scenarios, parameter estimates in the 100 fitting runs followed a smooth distribution not indicating the convergence of the runs towards different local minima (Supplementary Fig. 6 A and B) ^11,31^. All code to run and analyse the mathematical model is available at https://github.com/GrawLab/MalariaReplication.

---

Supplementary Table 1: Gametocyte-related genes published in Painter et al. ^32^, Poran et al. 2017^33^, Lopez-Barragan et al. 2011^34^ and Jeninga et al.^35^ and their expression in end-dry season asymptomatic infections and clinical malaria cases published in Andrade et al. 2020 ^11^. Data is available upon request.

**Supplementary Table 2:** Parameter prior ranges.

| Parameter | Unit | Description | Prior range | Reference |
| --- | --- | --- | --- | --- |
| ***IPL*** | Parasites/ml | Initial parasite load | 100 - 1000 | (Mouwenda et al 2024)<sup>75</sup> |
| **$\sigma_{\max}$** | h<sup>-1</sup> | Maximal removal rates during acute phase | 0.01-0.15 | (Andrade et al 2020) <sup>11</sup> |
| **$\sigma_{\max2}$** | h<sup>-1</sup> | Maximal removal rates during chronic phase | 0.01-0.4 |  |
| **$\delta_G$** | h<sup>-1</sup> (*10<sup>-4</sup>) | Death rate immature gametocytes | 0.1 – 1 | (Cao et al. 2019) <sup>74</sup> |
| **$\delta_{G5}$** | h<sup>-1</sup> (*10<sup>-2</sup>) | Death rate mature gametocytes | 0.135 – 3.25 | (Cao et al 2019, Eichner et al 2001) <sup>42,74</sup> |
| ***m*** | h<sup>-1</sup> (*10<sup>-2</sup>) | Gametocyte maturation rate | 1.38 - 4.1 | (Cao et al 2019, Eichner et al 2001) <sup>42,74</sup> |
| ***f*** | h<sup>-1</sup> (*10<sup>-2</sup>) | Commitment rate | 0.1 - 1.0 | (Cao et al 2019, Eichner et al 2001) <sup>42,74</sup> |
| ***f*<sub>2</sub>, *f*<sub>3</sub>** | h<sup>-1</sup> (*10<sup>-2</sup>)h – 1 (* 10 – 2) | Modified commitment rate<br>*Modified commitment rate* | 0.1 - 1.0 0.1 – 1.0 | (Cao et al 2019, Eichner et al 2001) <sup>42,74</sup> |

**Supplementary Table 3:**
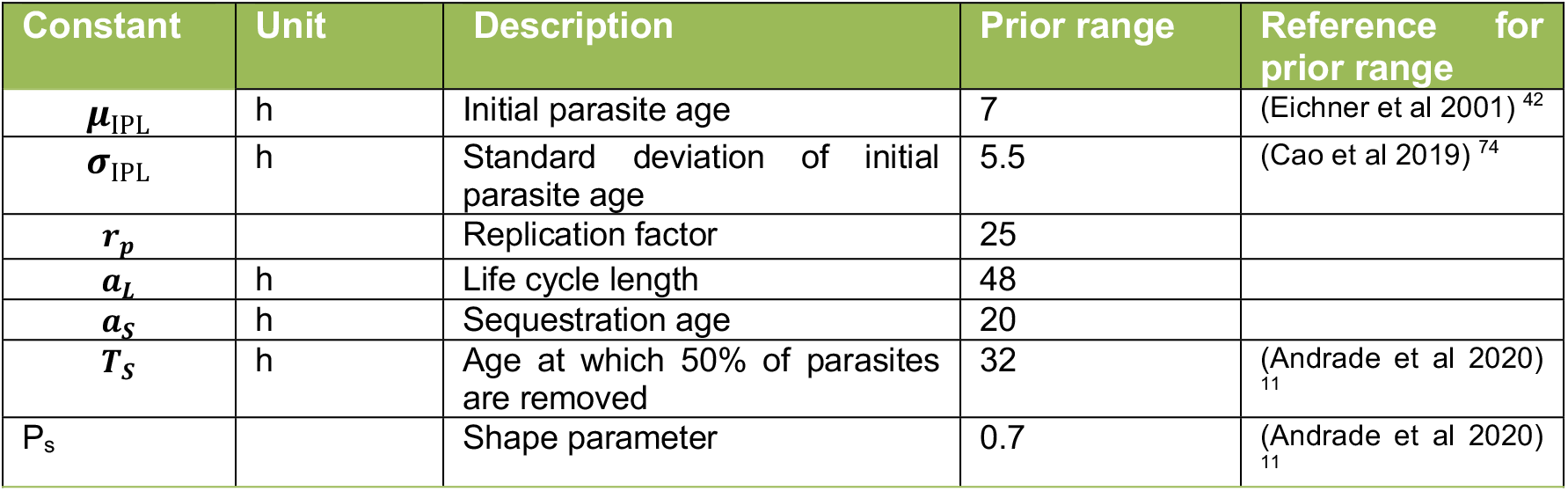
Model constants.

**Supplementary Table 4:** 89% CI of posterior parameter distribution.

| Parameter | Unit | Scenario 1 | Scenario 2 | Scenario 3 |
| --- | --- | --- | --- | --- |
| $IPL$ | Parasites/ml | 264 - 1020 | 311 - 1060 | 276 - 1010 |
| $\sigma_{max}$ | h<sup>-1</sup> | 0.128 - 0.142 | 0.126 - 0.142 | 0.128 - 0.141 |
| $\sigma_{max2}$ | h<sup>-1</sup> (*10<sup>-2</sup>) | 5.46 - 6.91 | 5.49 - 723 | 5.52 - 6.89 |
| $\delta_G$ | h<sup>-1</sup> (*10<sup>-4</sup>) | 0.273- 0.987 | 0.215 - 0.91 | 0.285 - 0.974 |
| $\delta_{G5}$ | h<sup>-1</sup> (*10<sup>-2</sup>) | 1.74 - 3.34 | 2.15 - 3.32 | 1.85 - 3.37 |
| $m$ | h<sup>-1</sup> (*10<sup>-2</sup>) | 3.47 - 5.46 | 3.08 - 5.45 | 2.1 - 5.21 |
| $f$ | h<sup>-1</sup> (*10<sup>-2</sup>) | 0.11 - 0.404 | 0.104 - 0.376 | 0.193 - 0.896 |
| $f_2, f_3$ | h<sup>-1</sup> (*10<sup>-2</sup>) | | 0.348 - 1.01 | 0.109 - 0.324 |

## Acknowledgments

We thank the residents of Kalifabougou, Mali and Nangonera, Uganda for participating in the studies. We acknowledge the support of the Flow Cytometry Core Facility of DKFZ, the Heidelberg University Metabolomics Core Technology Platform, and the Clinical Lipidomics Unit of Mainz University Medical Center. We thank Matthias Marti and Lauriane Sollelis at University of Glasgow for help and expertise with in vitro induction of parasite sexual commitment and gametocytogenesis. This work was supported by the European Research Council (ERC) under the European Union’s Horizon 2020 research and innovation programme (grant agreement No 759534), and the German Center for Infection research (DZIF). The Malian cohort study was supported by the Division of Intramural Research, National Institute of Allergy and Infectious Diseases of the National Institutes of Health; and the Ugandan study was supported by the National Institutes of Health as part of the International Centers of Excellence in Malaria Research program. Chica and Heinz-Schaller Foundation supported Frederik Graw.

## Data Availability

The RNA-seq data analyzed during the current study are available through (1) NCBI’s Gene Expression Omnibus (http://www.ncbi.nlm.nih.gov/geo/) under GEO Series accession number GSE148125 (Andrade et al. 2020), GSE72695 (Painter et al. 2018), GSE180985 (Jeninga et al. 2023); (2) GanBank with accession number SRP009370 (Lopez-Barrangan et al. 2011); and (3) NCBI Sequence Read Archive under accession code SRP116718 (Poran et al.).

---

**Supplementary Figure 1.**
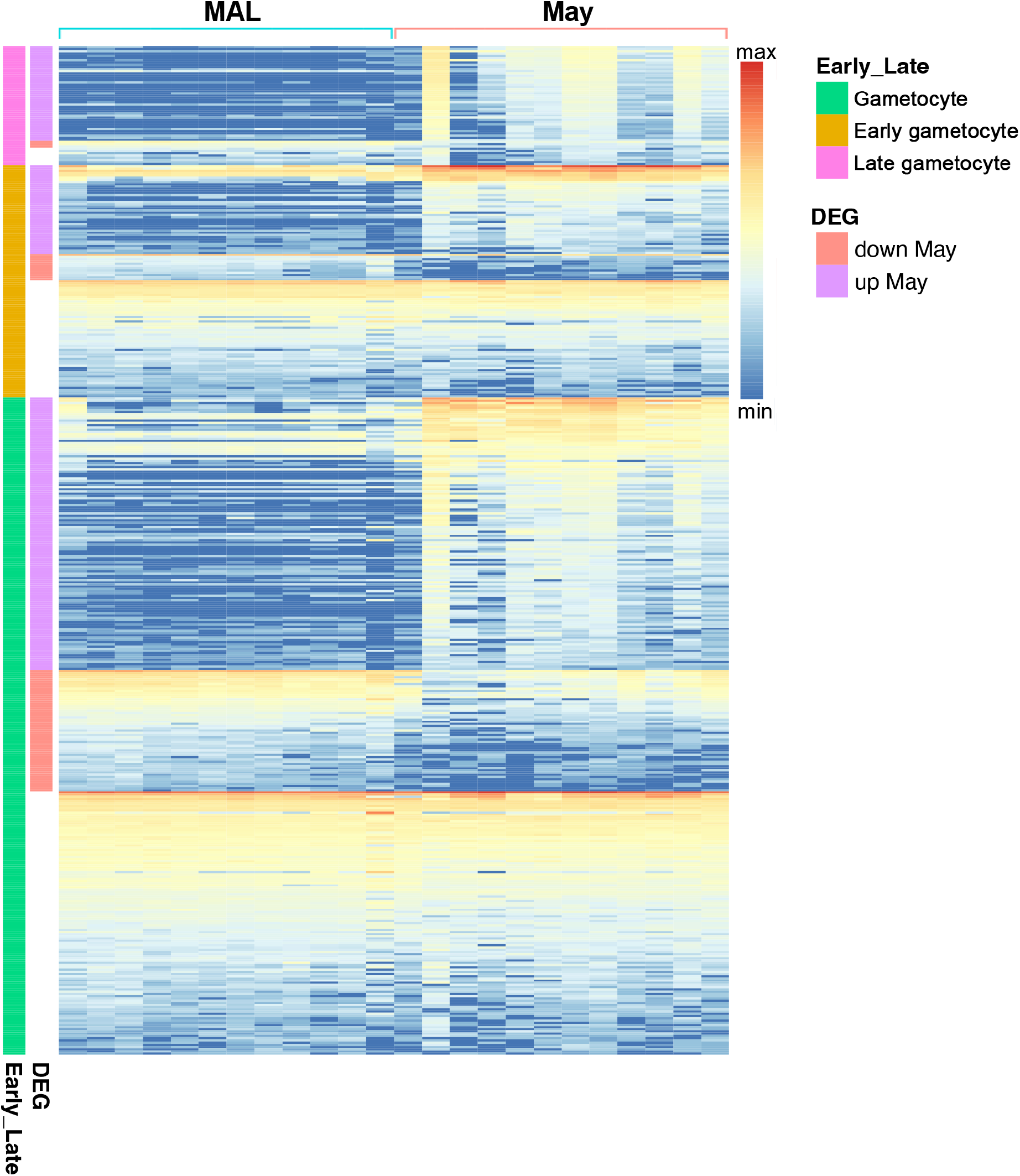
Heatmap showing normalized reads of 551 gametocyte specific genes of *P. falciparum* (rows) for each subject (columns) collected at the end of the dry season (May) and at the first clinical malaria case (MAL) in the ensuing transmission season.

**Supplementary Figure 2.**
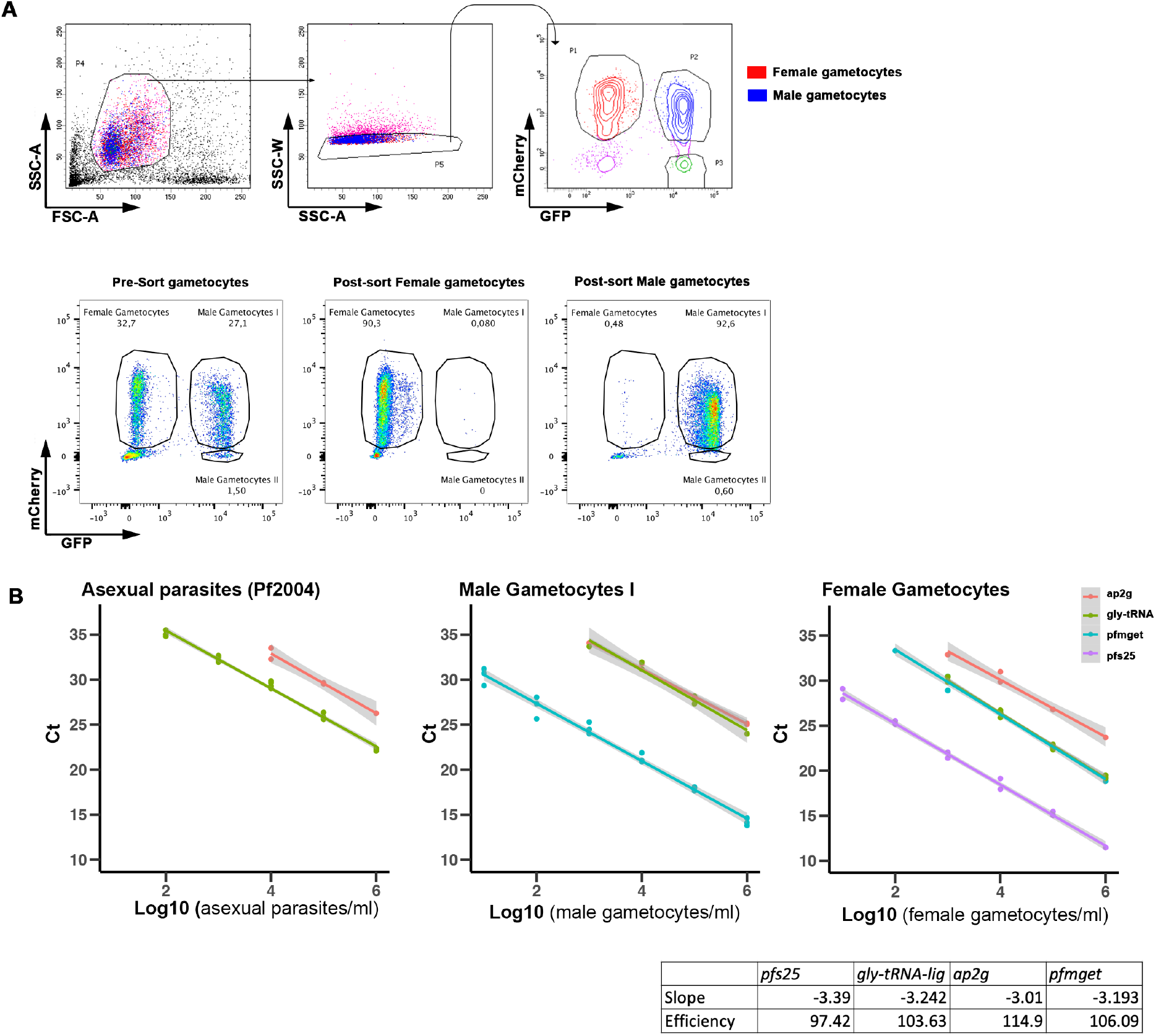
**A** Flow cytometry gating strategy to identify and sort PfDynGFP/ PfP47 gametocytes by mCherry and GFP expression into three populations (female: mCherry-positive, male I: mCherry-GFP-positive, male II: GFP-positive).**B** Pre-sort gametocyte distribution, and post-sorting purity verification of female and male gametocytes. **C** Serial dilutions of cDNA of asexual parasites and post-sort male and female gametocytes. Serial dilutions were used as standard curves to quantify total parasites, male and female gametocytes from q-RTPCR data of *pfs25, pfmget*, Glycine-tRNA ligase and to determine primer efficiency. Primer efficiency is shown on the figure. Primer efficiency of glycine-tRNA-ligase primers was determined based on asexual parasites, *pfs25* primer-efficiency based on female gametocytes and *ap2g* and *pfmget* in male gametocytes.

**Supplementary Figure 3.**
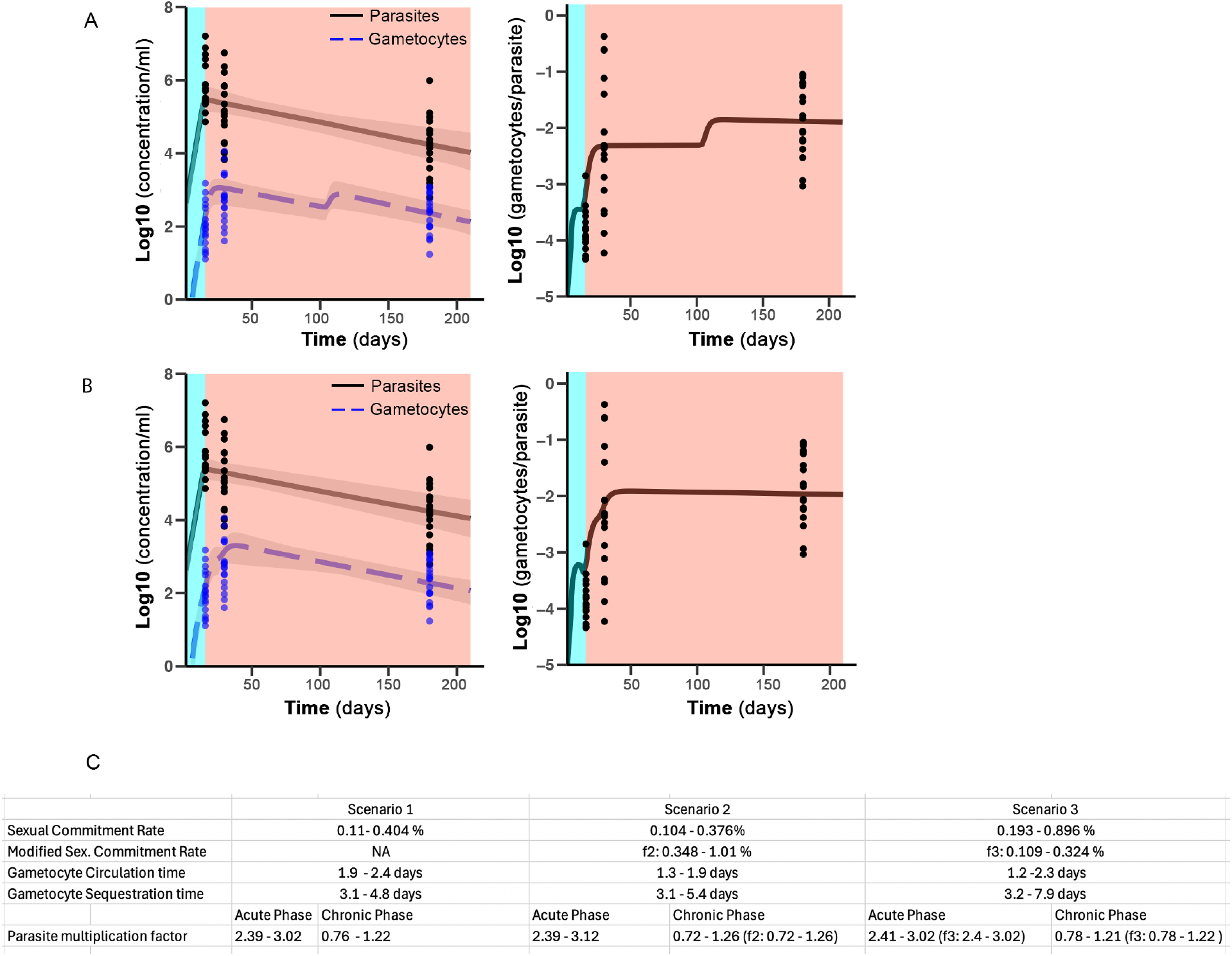
**A** Model prediction of development of parasitemia and gametocytemia (left) and gametocyte/parasite ratio (right) with infection progression in scenario 2. Average prediction is shown, grey shaded indicate the minimum and maximum estimate of parasitemia and gametocytemia based on posterior parameter distribution. Parasitemia and gametocytemia are shown averaged over a cycle length (48 h). Lines indicate model prediction, measurements for gametocytemia of MAL (d14), Jan (d28) and May (d180) are shown. **B** Development of parasitemia, gametocytemia and gametocyte/parasite ratio in scenario 3. Parasitemia and gametocytemia are shown averaged over a cycle length (48 h). Lines indicate model prediction, measurements for gametocytemia of MAL (d14), Jan (d28) and May (d180) are shown. **C** Biological parameter estimates of *P. falciparum* infection shown as 89% credibility interval of the posterior distributions in the three modelled scenarios.

**Supplementary Figure 4.**
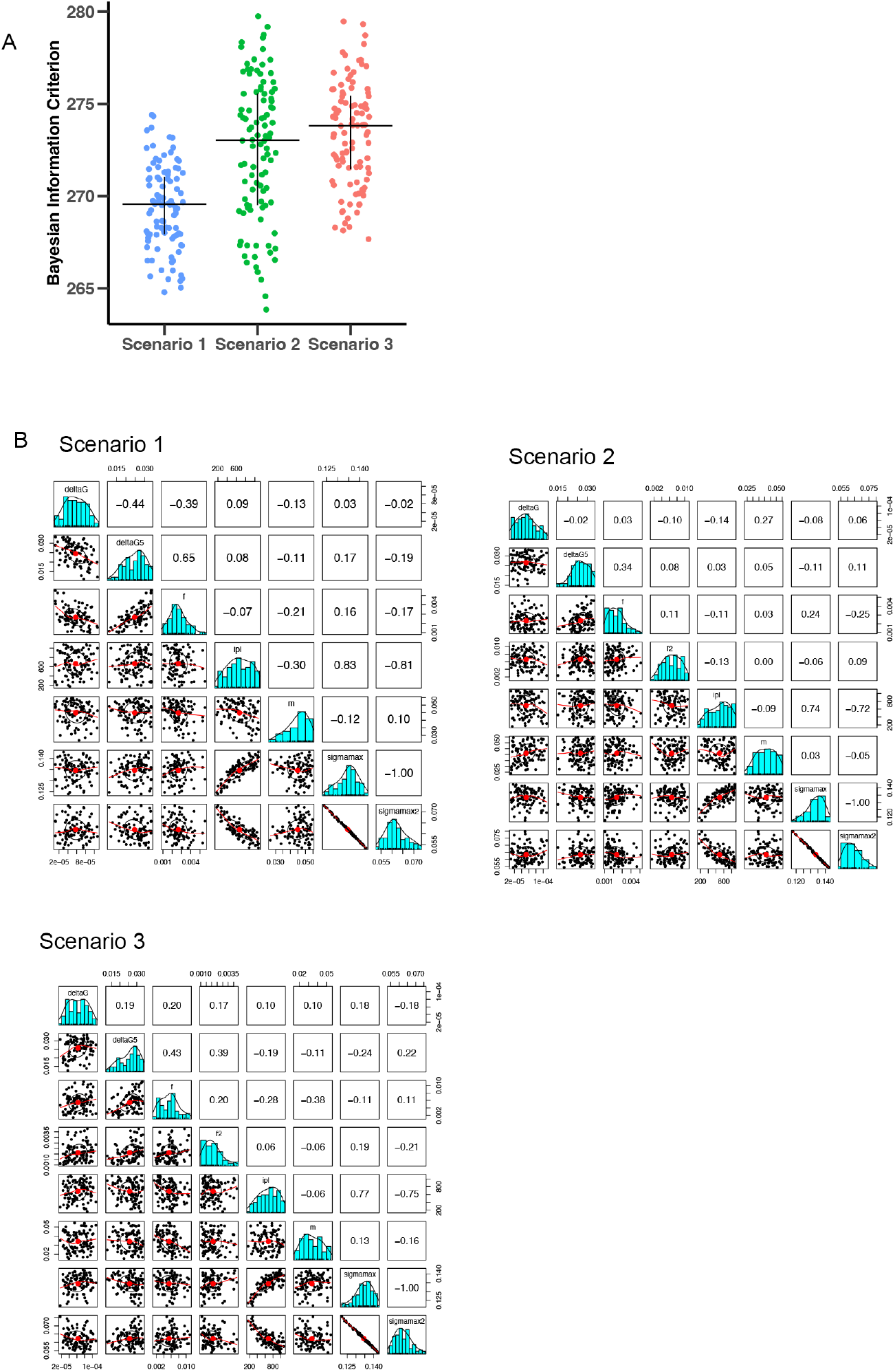
**A** Bayesian information criterion for the three scenarios calculated based on the parameter estimates in the 10^th^ pyABC generation after which the algorithm was stopped. **B** Correlation between parameter estimates in the final pyABC generation in the three scenarios. Posterior parameter distributions are shown on the diagonal, with the plots in the bottom triangular matrix showing scatterplots of each parameter combination, and numbers in the top triangular matrix indicating Pearson’s correlation coefficient of individual parameter combinations.

**Supplementary Figure 5.**
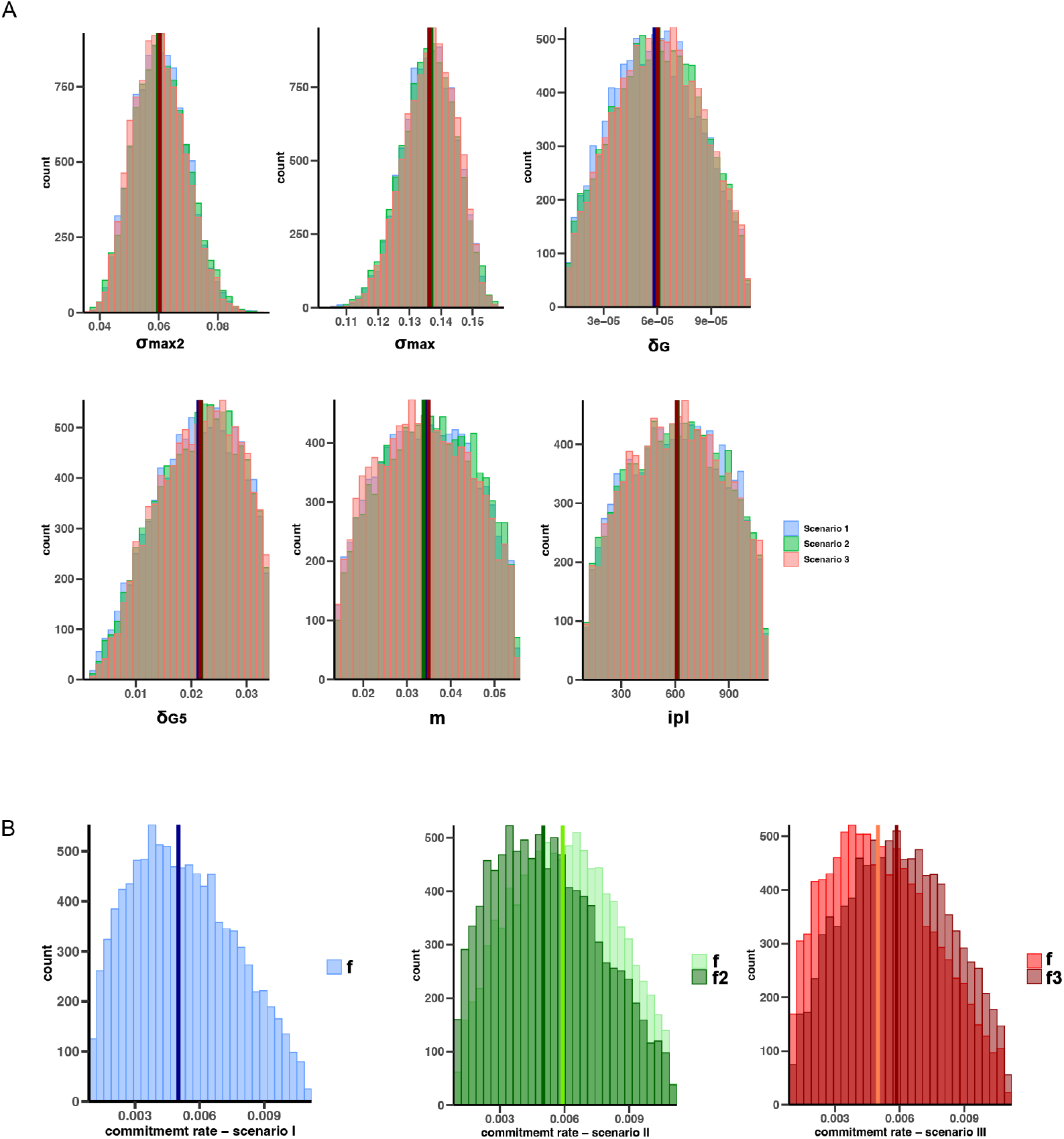
**A** Posterior parameter distributions of the three model scenarios of all parameters except the commitment rate. Histogram of parameter estimates from 100 pyABC fitting runs with each evaluated after the 10^th^ generation. Vertical lines indicate median parameter estimates. **B** Histogram of posterior distribution of the commitment rate in the three model scenarios, estimates from 100 pyABC fitting runs are shown with each evaluated after the 10^th^ generation.

**Supplementary Figure 6.**
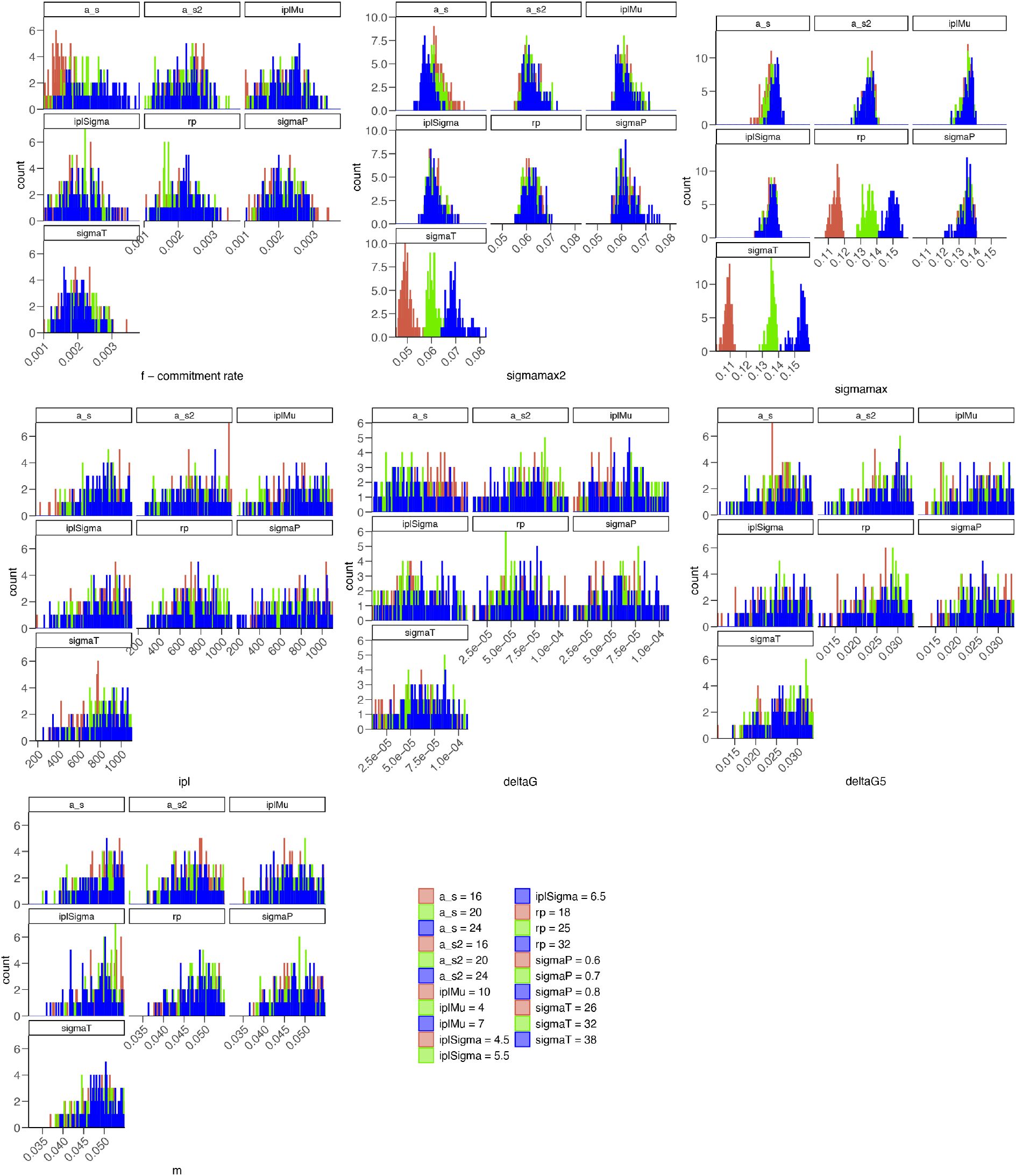
Sensitivity analysis. Histograms of posterior parameter distribution of model parameters in scenario I in response to variation of model constant. Parameters were fitted to a model with a high (blue), middle (green) and low (red) parameter value for each model constant. Each set of plots corresponds to one model parameter, with each small plot showing posterior distribution under variations of one model constant.

